# Structural Insights into the Trimeric Assembly of the HERC5 HECT Domain through SAXS Analysis

**DOI:** 10.1101/2025.03.22.644726

**Authors:** Cansu Deniz Tozkoparan Ceylan, Çağdaş Dağ

**Affiliations:** Nanofabrication and Nanocharacterization Center for Scientific and Technological Advanced Research (n^2^STAR), Koç University, İstanbul, Türkiye; Koç University Işbank Center for Infectious Diseases (KUISCID), Koç University, Istanbul, Türkiye

**Keywords:** HERC5, HECT domain, SAXS, ISGylation

## Abstract

HERC5 is an interferon-induced E3 ubiquitin ligase that mediates ISGylation, a critical post-translational modification involved in regulating antiviral immune responses. Its HECT (Homologous to the E6-AP Carboxyl Terminus) domain is essential for substrate recognition and the catalytic transfer of ubiquitin or ubiquitin-like proteins. Members of the HERC protein family are structurally defined by a C-terminal HECT domain and N-terminal RCC1-like (Regulator of Chromosome Condensation 1) domains. Despite its biological importance, the oligomerization behavior of the HERC5 HECT domain remains poorly understood. In this study, we examined the oligomeric state of the HERC5 HECT domain in solution using affinity and size-exclusion chromatography, followed by small-angle X-ray scattering (SAXS) at a protein concentration of 2.0 mg/ml and temperature of 10°C. SAXS data analysis revealed that the HERC5 HECT domain predominantly adopts a trimeric assembly. This conclusion is supported by Guinier and Kratky analyses, distance distribution functions, and ab initio shape reconstructions. These findings suggest that trimerization may play a regulatory role in the functional activity of HERC5, contributing to a deeper understanding of the structural mechanisms governing HECT-type E3 ligases.

## Introduction

The 15 kDa ubiquitin-like protein ISG15 (Interferon-Stimulated Gene 15) covalently attaches to target proteins through a process known as ISGylation. This modification plays a critical role in regulating antiviral responses within the mammalian innate immune system. ISGylation is mediated by a cascade of three enzymes: the E1 activating enzyme (Uba7), the E2 conjugating enzyme (UbcH8), and the E3 ligase enzyme (HERC5) [1,2].

E3 ubiquitin ligases are generally classified into four main families: homologous to E6AP C terminus (HECT), really interesting new genes (RING), RING-between-RINGs (RBR), and U-box-type ligases [3]. Among these, HECT-type E3 ligases are distinguished by a conserved catalytic domain of approximately 350 amino acids. This domain is composed of N-terminal and C-terminal lobes connected by a flexible linker and contains a catalytic cysteine residue (Cys994). The catalytic cysteine forms an intermediate thioester bond with ISG15 transferred from the E2 enzyme before the final transfer to substrate proteins [4–6]. The C-terminal lobe of the HECT domain mediates the catalytic transfer of ISG15, while the N-terminal lobe is primarily involved in substrate recognition and binding. Notably, HECT ligases can adopt different conformational states, existing as monomers or higher-order oligomers, which can significantly influence their enzymatic activity and the mechanism of ISG15 (or ubiquitin) transfer [4–6].

The E3 ligase enzyme involved in ISGylation, HERC5, interacts with the E2 conjugating enzyme UbcH8 via its HECT domain, thereby facilitating the transfer of ISG15 to target proteins. HERC5 is a 117 kDa antiviral protein induced by interferon-alpha/beta (IFN-α/β) signaling. HERC5 possesses the unique ability to associate with polyribosomes and mediate co-translational ISGylation, particularly in virus-infected host cells, thereby modifying newly synthesized proteins as part of the antiviral defense response. [4,7]. HERC5 functions as a positive regulator of innate immunity against HIV-1 by modifying viral Gag proteins with ISG15, which disrupts viral assembly and budding [8]. Despite its biological importance, the molecular mechanism of HERC5—particularly the structural and dynamic properties of its HECT domain—remains poorly understood.

There is currently no universal consensus regarding the precise boundaries of the HECT domain within HECT-type E3 ligases. While the core catalytic HECT domain is generally recognized to consist of approximately 350 amino acids encompassing the conserved N-and C-lobes and the catalytic cysteine, some studies include an additional N-terminal α-helix preceding the canonical HECT fold as part of the domain, whereas others exclude it (Figure 1). In our previous work, through sequence alignment and structural comparison with the well-characterized E6AP HECT domain, we demonstrated that the HERC5

**Figure 1.**
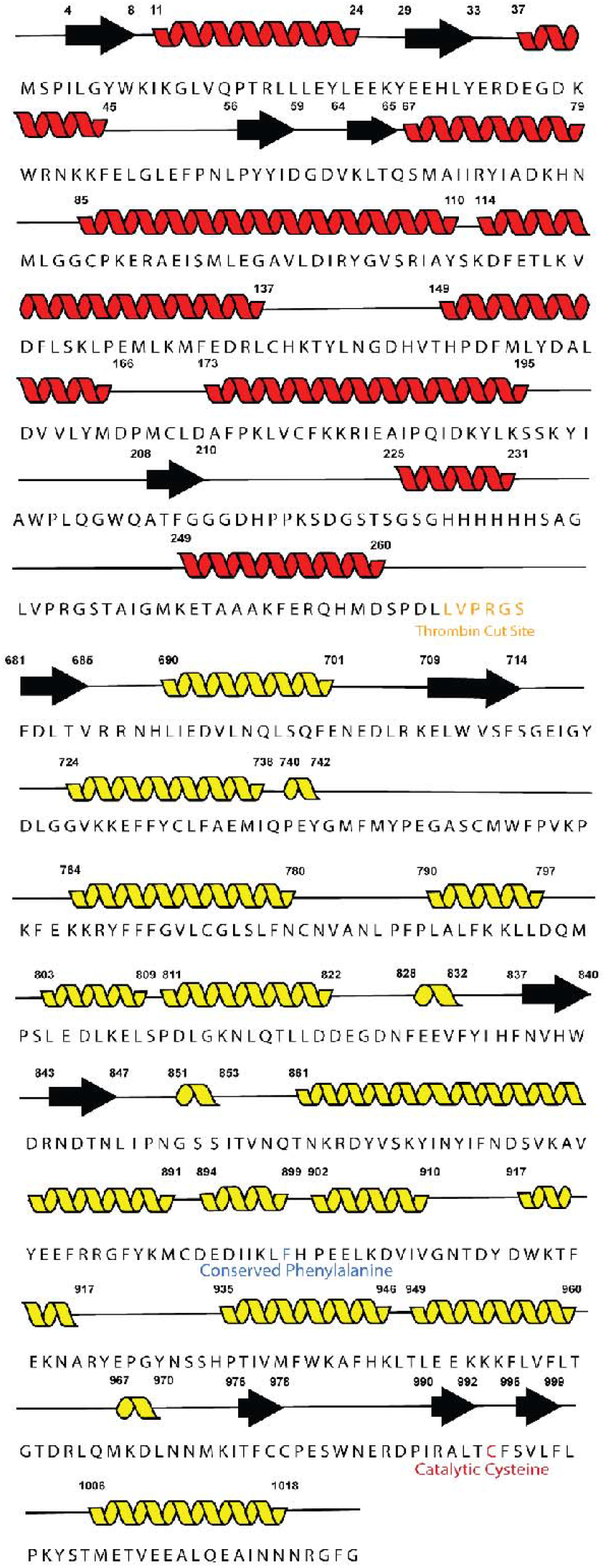
Representation of the GST-HERC5 HECT domain construct showing the amino acid sequence and predicted secondary structure. GST is shown in red, the HERC5 HECT domain in yellow, the thrombin cleavage site in orange, the conserved phenylalanine residue in blue, and the catalytic cysteine (Cys994) in dark red.

HECT domain shows a similar domain architecture [10]. Specifically, our analysis identified the HERC5 HECT domain as spanning residues 681 to 1024, corresponding to a length of 344 amino acids and a molecular mass of approximately 40.5 kDa (Figure 1). This definition aligns with the functional and structural core necessary for ISG15 transfer, while acknowledging potential variability in domain annotations across different studies.

Small-angle X-ray scattering (SAXS) is a versatile structural technique widely applied in the study of biological macromolecules. In protein research, SAXS provides valuable information on particle size, shape, and conformational states in solution [9]. SAXS experiments yield two main data components: the scattering vector (q) and the scattering intensity (I(q)). These data are typically analyzed using the Fourier transform, and results are commonly displayed as log–log plots where the logarithm of I(q) is plotted against the logarithm of q.

In these log–log plots, three characteristic regions can be identified: the Guinier region at low q, a central region, and the Porod region at high q. The Guinier plot, which graphs ln I(q) versus q^2^ at low q values, provides the radius of gyration (Rg), a measure of the overall particle size. A linear Guinier region indicates high-quality data without aggregation or interparticle effects. The Kratky plot (q^2^I(q) versus q) helps assess whether the protein is folded, partially folded, or intrinsically disordered. Porod analysis focuses on the high-q region and reveals information about the particle’s surface properties, such as whether it is smooth or rough. Additionally, the pair-distance distribution function P(r) offers detailed insights into the shape and flexibility of particles, distinguishing between globular, elongated, multi-domain, or unfolded conformations in solution.

In this study, we investigated the structural organization and conformational properties of the C-terminal HECT domain of HERC5 in solution using small-angle X-ray scattering (SAXS). The SAXS data provide valuable insights into the overall shape, molecular dimensions, flexibility, and conformational variability of the HERC5 HECT domain. When considered in the context of existing literature, these findings offer new perspectives on the oligomeric behavior of HERC5 in solution and its potential influence on structural dynamics. Importantly, our results contribute to a deeper understanding of the ISG15 transfer mechanism mediated by HERC5, and lay the groundwork for future biotechnological and pharmacological strategies targeting this antiviral E3 ligase.

## Material & Method

### Chemicals

Lysogeny broth (LB), kanamisin, chloramphenicol, isopropyl-1-thio-β-d-galactopyranoside (IPTG), TRIS (tris(hydroxymethyl)aminomethane), sodium chloride (NaCl), dithiothreitol (DTT), Glutathione-s-transferase, and all other common chemicals were obtained from Sigma (USA).

### Protein expression and Purification

The HECT domain of HERC5 (residues 702-1024) with a glutathione-S-transferase (GST) fusion protein was expressed in *Escherichia coli* BL21 (DE3) (Thermo Fisher Scientific, USA) and purified as described [10]. The eluted free HERC5 HECT protein fractions were pooled together, concentrated to about 5 ml by 30K centrifugal ultrafilters. Size-exclusion chromatography was carried out in the ÄKTA™ ^™^ start protein purification system chromatography system using the Superdex S75 16/60 Column. The column was equilibrated with an equilibration buffer (20 mM Tris-base,150 mM NaCl, and 1 mM DTT pH: 7.4) and samples were loaded onto the column. The elution fractions were collected by SEC buffer (20 mM TRIS, 150 mM NaCl and 1 mM DTT pH: 7.4).

### Small Angle X-Ray Scattering (SAXS) data collection, processing, and modeling

Small-angle X-Ray scattering (SAXS) experiments were performed to determine native state of HERC5-HECT in solution. The SAXS experiment was performed using the SAXSpace instrument (Anton Paar 5.0), with a ‘Multi Cuvette Holder’, ‘Heated/Cooled Sample Holder’, and a ‘Beam Stopper’ utilizing a 2 mm Ni. Furthermore, the sample-to-detector distance (SDD) is set to 1550 nm, and the 6 frames are set to last 1200 seconds each. Then, SAXS analysis was initiated at 2.0 mg/ml at 10 °C using an 80 μl of protein sample. The gathered two-dimensional scattering data was processed using various steps, including Zero Point 2D by Moments, Masking, Q Transformation, Transmission, Data Reduction, and Standard Operations 1D in SAXSanalysis™ software [11]. The generated ‘.dat’ file was subsequently uploaded to the PRIMUS package within ATSAS 4.0 to compute the log-log plot, Guinier plot, Kratky plot, pair-wise distance distribution function P(R), radius of gyration (Rg), scattering intensity I(0), maximum particle size in solution (Dmax), and Porod volume (Vp) values. The graphical data revealed that the protein existed in a trimeric form, prompting the use of the DAMMIF tool to produce low-resolution initial models of protein structures by analyzing. The obtained model was overlayed with HERC5 HECT protein (Q9UII4) Alphafold structure.As no experimental structure is currently available for HERC5, we used the AlphaFold-predicted structure of the HERC5 HECT domain to interpret our SAXS data. The AlphaFold model represents the monomeric form of the domain. To construct a trimeric model consistent with our SAXS findings, we aligned three copies of the HERC5 HECT monomer onto the trimeric crystal structure of the structurally related E6AP HECT domain. This composite model provided a good fit to the SAXS envelope, supporting the proposed trimeric assembly of HERC5. The overlay was visualized by PyMol software.

## Results and Discussion

Small-angle X-ray scattering (SAXS) experiments were performed to investigate the structural features and solution behavior of the HERC5 HECT domain. These measurements provided insights into the overall morphology, conformational flexibility, and potential protein–protein interactions of HERC5 in solution. SAXS data were collected at a protein concentration of 2.0 mg/mL, and the derived structural parameters are summarized in Table 1 and Table 2.

**Table 1.** SAXS Instrument parameters of HERC5 HECT.

|  |  |
| --- | --- |
| Instrument | SAXSpoint 5.0 |
| s range (nm<sup>-1</sup>) | 0.01 nm<sup>-1</sup> to 49.3 nm<sup>-1</sup> |
| Exposure time (min) | 120 |
| Temperature (C°) | 10 |
| Sample volume (μl) | 80 |
| Sample concentration (mg/ml) | 2.0 |

**Table 2.**
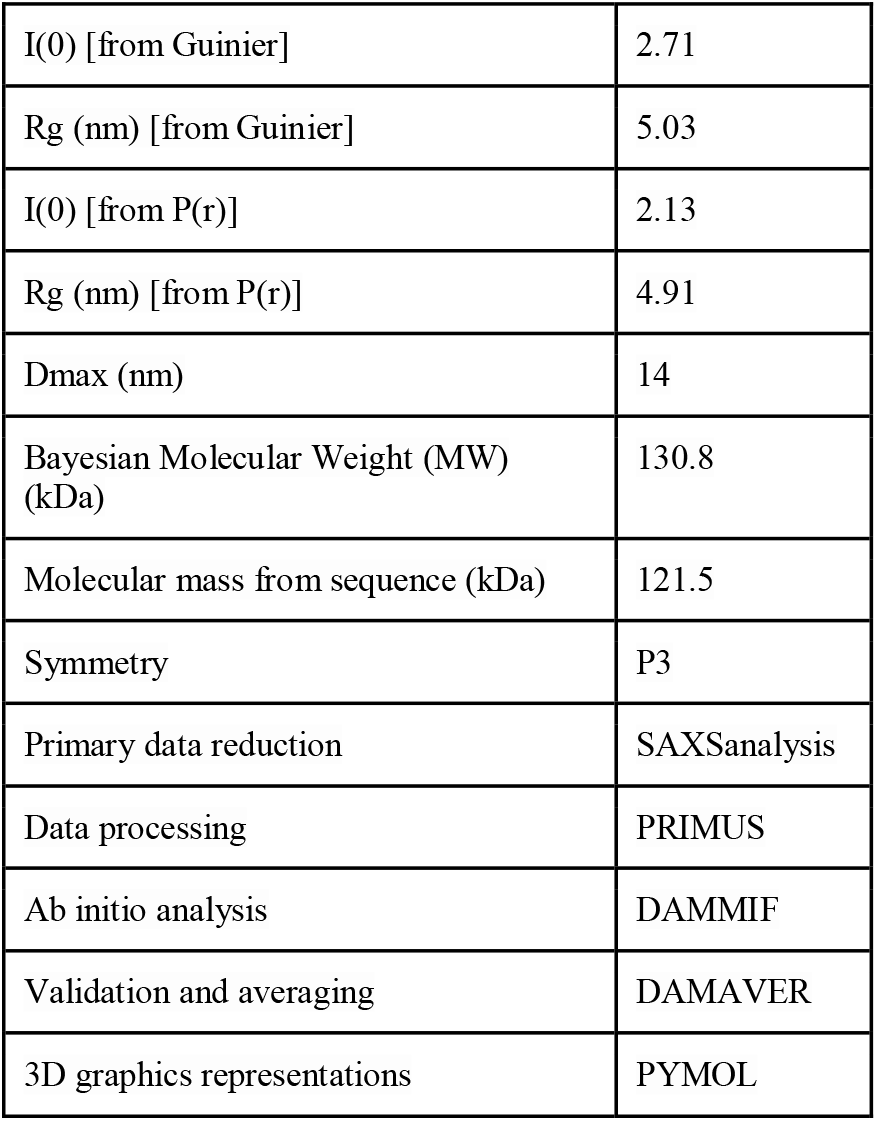
SAXS parameters of HERC5 HECT.

The Rg values derived from Guinier analysis are presented in Table 2. The radius of gyration (Rg), calculated using Guinier analysis, was determined as 5.03 nm, closely matching the value obtained from the pair-distance distribution function (P(r)), 4.91 nm. The similarity of Rg values from both methods underscores the consistency and reliability of the SAXS data. The maximum particle dimension (D_max) derived from the P(r) function was found to be 14 nm, supporting the presence of a compact oligomeric structure rather than an elongated or unfolded conformation.The molecular mass of the HERC5 HECT trimer, estimated by Bayesian inference, was 130.8 kDa. This closely agrees with the theoretical molecular mass calculated from the amino acid sequence (121.5 kDa for a trimer), further reinforcing the conclusion of a predominantly trimeric oligomerization state under the conditions tested. Consistent Rg and I(0) values were also obtained from the pair-distance distribution function [P(r)], demonstrating good agreement between reciprocal and real space data and confirming the reliability of the SAXS measurements [12]. Figure 2 presents comprehensive SAXS data analysis for the HERC5 HECT domain in solution. Figure 2A displays the SAXS scattering profile as log-intensity versus scattering vector (q), demonstrating a typical scattering curve of a folded protein with well-defined scattering features. Guinier analysis reveals a distinct linear region at low q^2^, indicative of high-quality data without significant aggregation or polydispersity, confirming suitability for precise radius of gyration (Rg) determination. Figure 2B shows the fitting of the SAXS experimental data with theoretical scattering calculated by GNOM. The high-quality fit between experimental and theoretical curves supports the accuracy of derived structural parameters. Figure 2C presents the pair-distance distribution function [P(r)], which indicates a bell-shaped curve typical of globular proteins, reflecting a clearly defined maximum dimension (D_max) and confirming a compact, non-extended conformation of the protein in solution. The observed asymmetry in the P(r) distribution likely reflects the inherent domain organization of the HERC5 HECT domain, which contains an N-lobe and a C-lobe connected by a flexible linker. Such two-domain architectures typically produce skewed P(r) curves due to a broader range of intramolecular distances, and the result is consistent with the known structural arrangement of HECT E3 ligases. The P(r) function is consistent with the presence of a distinct oligomeric state rather than an unfolded or elongated structure. Panel D provides the Kratky plot, illustrating a clear peak at low-to-intermediate q-values followed by convergence at higher q-values. This pattern is characteristic of well-folded, globular macromolecules with limited flexibility or disorder. The absence of a pronounced elevation in the higher q-region further confirms that the HERC5 HECT domain is predominantly compact and well-structured, supporting the conclusion of a stable oligomeric assembly in solution [13,14]. Collectively, these results strongly support the presence of a well-defined, globular oligomeric structure of the HERC5 HECT domain, consistent with a trimeric assembly as suggested by complementary size-exclusion chromatography analyses [15,16].

**Figure 2.**
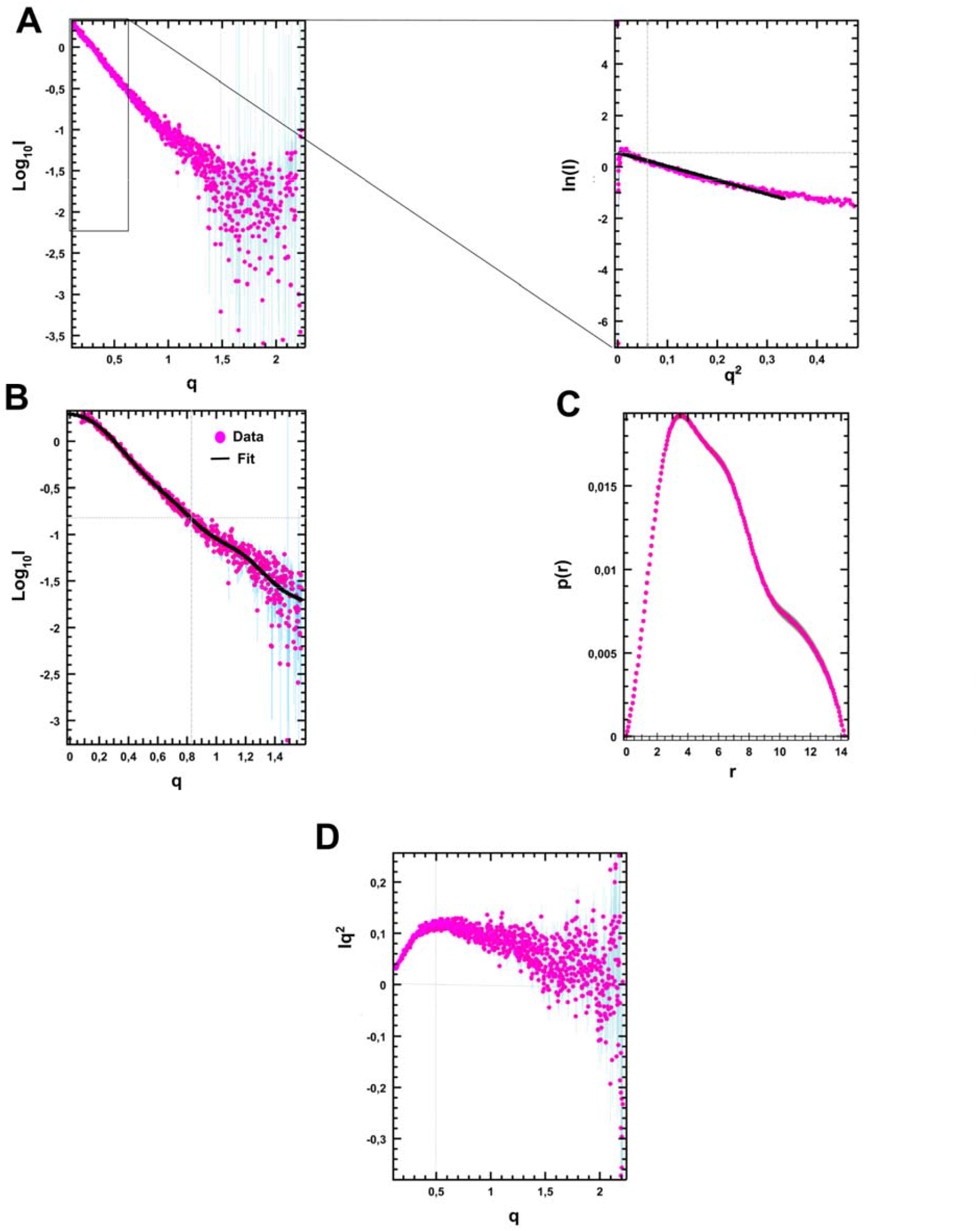
Figure 2. Small-angle X-ray scattering (SAXS) analysis of the HERC5 HECT domain. A. Log–log plot of scattering intensity logI(q) versus scattering vector q (nm□^1^). B. Fit of the experimental SAXS data to the GNOM-calculated curve as a function of q (nm□^1^). C. Pair-distance distribution function P(r) plotted against real-space distance r (nm). D. Kratky plot of q2I(q) versus q (nm□^1^), indicating particle compactness and flexibility.

The molecular weight estimates presented in Table 2, which are almost three times higher than the theoretical molecular weight of the protein suggest that the protein forms a trimer in solution. Low-resolution bead envelopes for the HERC5 HECT protein were constructed using the DAMMIF program [14, 15, 17]. The AlphaFold structure of the HERC5 HECT protein (Q9UII4) was superimposed with the DAMAVER model, which averaged the 21 DAMMIF generated images. The ab initio model was fitted using PyMOL software (Figure 3). To evaluate the quality of the SAXS-derived models and their agreement with experimental data, several fit metrics were calculated. The goodness-of-fit (χ^2^) value between the experimental SAXS profile and the theoretical scattering curve generated by GNOM from the pair-distance distribution function P(r) was 1.12, indicating a high-quality fit. The fit between the experimental data and the averaged ab initio model obtained using DAMAVER yielded a χ^2^ value of 1.34, consistent with well-resolved low-resolution reconstructions. To further assess structural agreement, the AlphaFold-predicted monomeric model of the HERC5 HECT domain was superimposed onto the SAXS-derived ab initio envelope of the trimeric assembly. The spatial correspondence was quantified using the normalized spatial discrepancy calculated by SUPCOMB, resulting in a value of 0.87, supporting a reasonable alignment between the AlphaFold-derived model and the SAXS-based shape. These values together indicate reliable consistency between experimental SAXS data and computational modeling.

**Figure 3.**
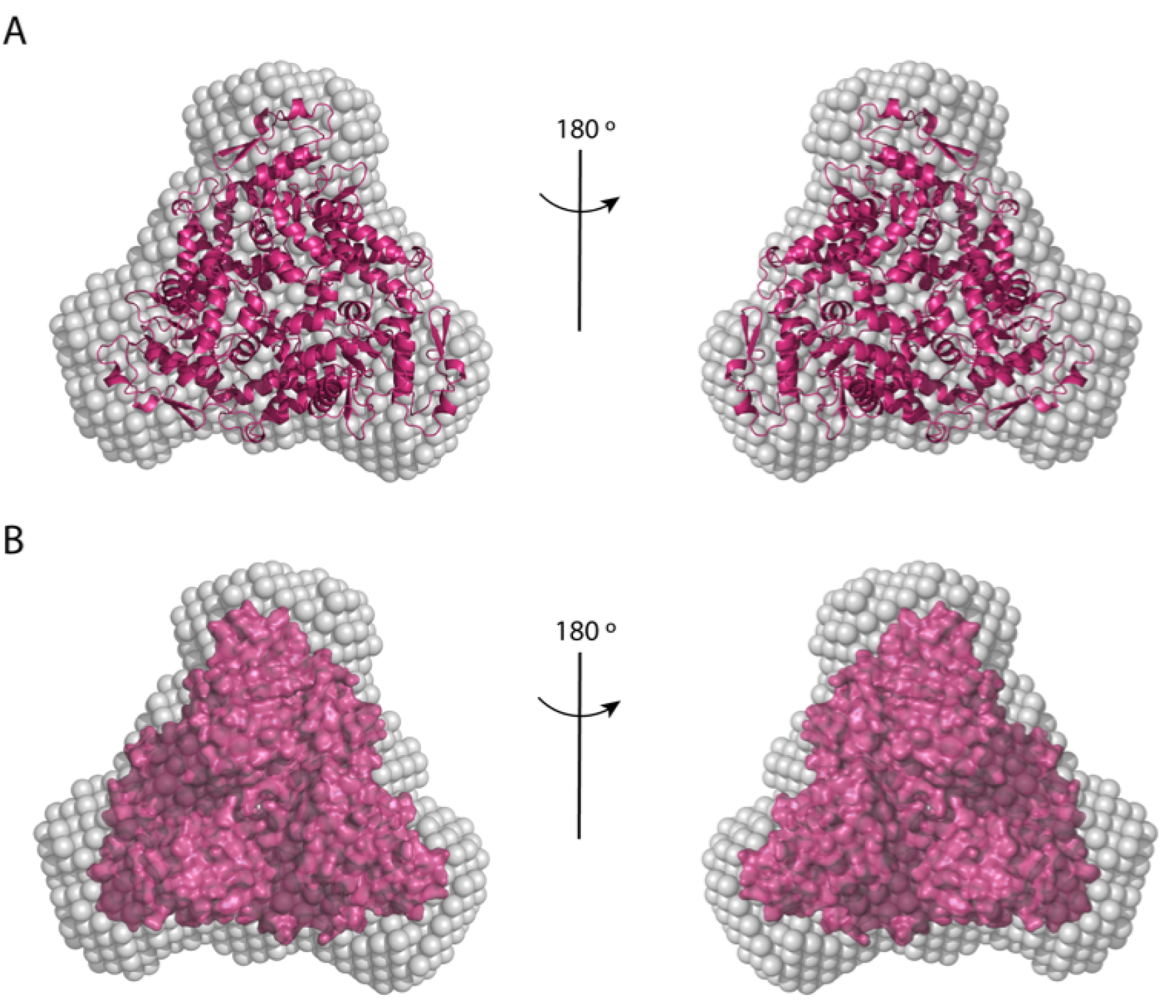
Cartoon (A) and surface (B) representations of the trimeric model of HERC5 HECT, generated by aligning three AlphaFold-predicted monomers (AlphaFold ID: Q9UII4) onto the trimeric crystal structure of the E6AP HECT domain. The model is superimposed on the SAXS ab initio envelope reconstructed using DAMMIF with P3 symmetry.

Structural analysis of the HERC5 HECT domain oligomerization interface identified several key interfacial residues likely responsible for stabilizing the observed trimeric assembly (Figure 4). The model, generated by aligning three AlphaFold-predicted HERC5 monomers onto the known trimeric crystal structure of the E6AP HECT domain, reveals that residues Tyr720, Leu722, Lys807, Asp822, and Gln818 participate in close inter-subunit contacts through potential hydrophobic interactions and electrostatic bonds. Notably, the catalytic cysteine (Cys994) is positioned away from the oligomerization interface, suggesting that oligomerization does not directly obstruct catalytic site accessibility.The obtained ab initio SAXS envelope was superimposed with the AlphaFold-predicted monomeric structure of the HERC5 HECT domain to evaluate structural compatibility and potential oligomerization states. Unlike several other HECT-type ligases such as WWP1 [18], Smurf2 [19], HUWE1 [20], and Nedd4-1 [21], which do not show trimeric forms, the closely related E6AP ligase has been reported to uniquely adopt a trimeric configuration [22]. Although E6AP was initially hypothesized to function primarily as a monomer, Huang et al. (1999) provided early structural evidence suggesting a potential trimeric arrangement formed by crystal packing interactions [22]. Subsequent kinetic and biophysical studies have demonstrated that E6AP indeed requires oligomerization, specifically trimerization, for efficient assembly of polyubiquitin degradation signals [23]. Collectively, these observations lend support to our SAXS-based evidence that the HERC5 HECT domain similarly adopts a trimeric oligomerization state, reinforcing the functional relevance of oligomerization among HECT-type E3 ubiquitin ligases.

**Figure 4.**
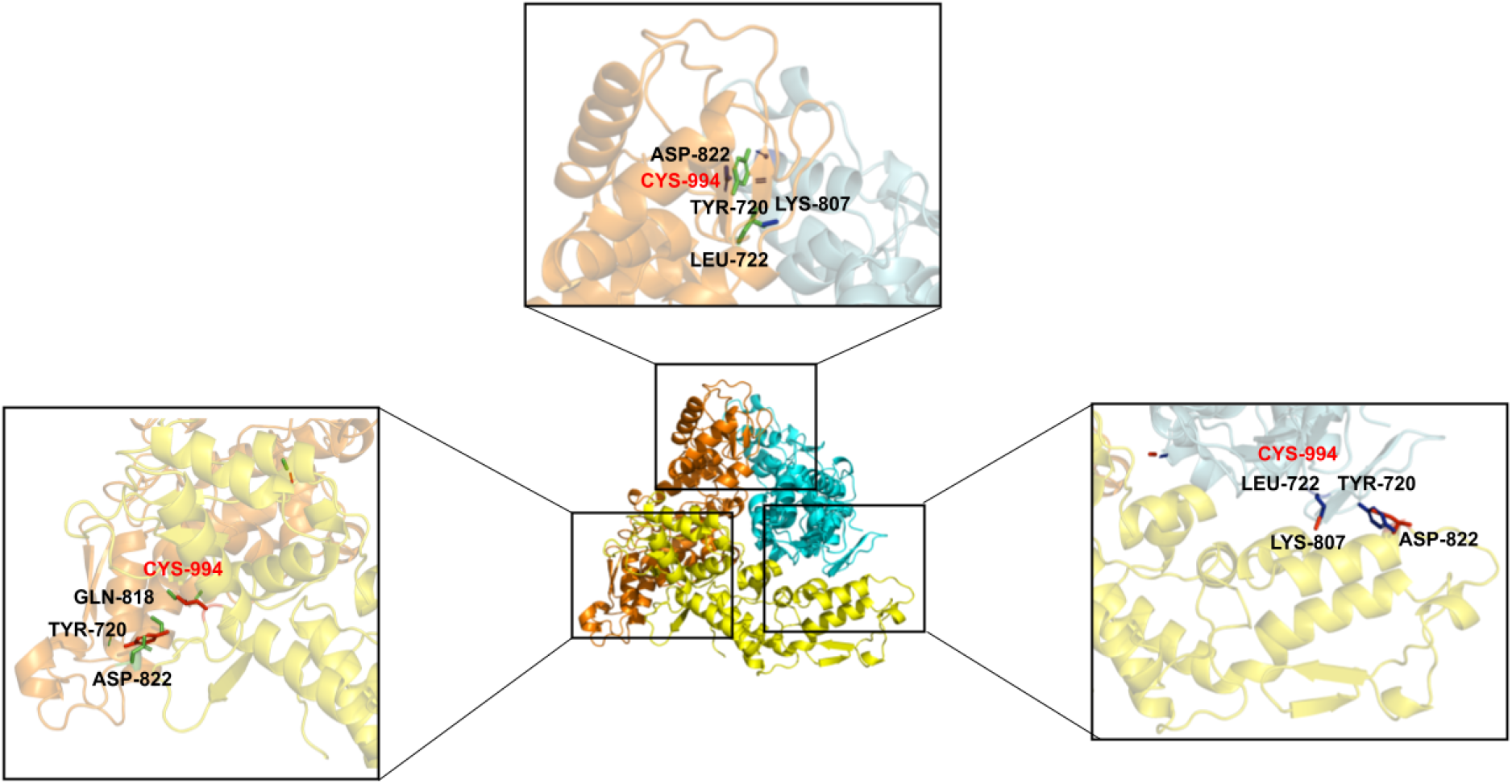
Interfacial side-chains contacts in the trimeric HERC5 HECT domain.

Previous studies have highlighted the significance of residue Phe727 in the oligomerization of E6AP, specifically through its contribution to hydrophobic interactions between subunits, thus stabilizing the trimeric assembly. Mutations at this critical position have been shown to impair trimer formation and enzymatic activity, linking such alterations to neurodevelopmental disorders, including Angelman Syndrome. Notably, the HPV E6 viral protein enhances the degradation of tumor suppressors such as p53 by promoting E6AP oligomerization through its homodimerization capability, consequently increasing catalytic activity. Further kinetic analyses indicated that despite mutations affecting trimer stability, E6AP retains its ability to bind the E2 conjugating enzyme, and the active-site cysteine (Cys820) within the HECT domain remains competent in forming the intermediate ubiquitin-thioester complex essential for ubiquitin transfer [23].

Expanding on this mechanistic insight, Ronchi et al. (2017) combined in silico modeling with kinetic studies to demonstrate that E6AP possesses two functionally distinct E2 binding sites, challenging the conventional single-site transthiolation model. Their findings proposed that these separate E2-binding sites, located on different subunits within the E6AP trimer, cooperatively coordinate polyubiquitin chain assembly [24]. Consistently, earlier experimental evidence from Ronchi et al. (2014) established that the catalytically active form of E6AP is indeed a trimer, which is indispensable for efficient polyubiquitin chain formation. Structural investigations by Takade et al. (2017), using high-speed atomic force microscopy (HS-AFM), revealed considerable conformational dynamics within the E6AP HECT domain, identifying distinct conformations including L-shaped, inverted T, and catalytic forms. Their analyses primarily detected monomeric and dimeric states, indicating the possibility of diverse oligomerization patterns within HECT domains [25].

Our present study demonstrates that the closely related HERC5 protein exhibits a trimeric oligomerization state of its HECT domain in solution, as supported by SAXS analyses indicating a molecular weight consistent with threefold the theoretical monomeric mass. This trimeric arrangement is further validated by bead-model fitting of the SAXS-derived envelope. Todaro et al. (2018) reported that another HECT-type ligase, NEDD4-2, similarly utilizes oligomerization to regulate its enzymatic activity, with residue Phe823 identified as crucial for stabilizing the trimeric assembly. Given the structural and functional similarity between NEDD4-2 Phe823 and E6AP Phe727, oligomerization appears to be a conserved mechanism regulating ubiquitin chain specificity, particularly influencing Lys63-linked polyubiquitination [26]. Recent structural insights provided by Hodáková et al. (2023) further extend our understanding by demonstrating that the HECT ligase UBR5 can adopt higher-order oligomeric states, specifically dimeric and tetrameric forms, identified through cryo-electron microscopy. Their findings revealed an intriguing cage-like tetrameric structure formed by face-to-face pairing of crescent-shaped dimers, positioning all catalytic HECT domains toward a central cavity, underscoring the broader relevance of oligomerization as a critical regulatory mechanism among HECT E3 ubiquitin ligases [27].

Oligomerization is increasingly recognized as a key regulatory mechanism among HECT-type E3 ligases, with functional implications for substrate binding, catalytic efficiency, and spatial organization. Previous studies have reported dimeric and tetrameric assemblies of HERC family proteins, including HERC6 and engineered constructs of HERC5, suggesting that multiple oligomeric states may coexist or be dynamically regulated under different physiological conditions. In our study, SAXS analysis of the HERC5 HECT domain revealed a trimeric assembly in solution, which has not been previously described for this protein. The structural modeling, supported by fitting to the ab initio SAXS envelope, indicates a well-ordered trimer that may provide a platform for cooperative interactions or increased local concentration of catalytic sites. Our SAXS-based structural analyses strongly suggest a stable trimeric oligomerization state for the HERC5 HECT domain. These findings align closely with prior reports on structurally related HECT-type E3 ligases such as E6AP and NEDD4-2, where oligomerization has been shown to be essential for polyubiquitin chain assembly and enzymatic activity [23,24,26]. For instance, oligomerization-dependent stabilization through conserved hydrophobic interactions, notably involving residues analogous to Phe727 in E6AP and Phe823 in NEDD4-2, has been demonstrated as critical for the catalytic function of these enzymes. Our identification of a structurally homologous residue (Phe899) within the HERC5 HECT domain, analogous to E6AP Phe727, further supports the functional relevance of trimer formation observed in our study.

The structural conservation across the HECT ligase family implies that oligomerization could be a general regulatory mechanism affecting both enzymatic activity and specificity toward substrates. Given the importance of HERC5 in antiviral immunity, particularly through ISGylation of nascent viral proteins, the trimeric conformation identified here might represent a biologically relevant active state, optimizing enzyme-substrate interactions in virus-infected cells. Therefore, targeting oligomerization interfaces, as previously suggested for E6AP and NEDD4-2, could present novel opportunities for therapeutic intervention aimed at modulating HERC5 activity in antiviral defense mechanisms.

## ACKNOWLEDGEMENTS

The authors acknowledge the use of the services and facilities of n2STAR-Koç University Nanofabrication and Nanocharacterization Center for Scientific and Technological Advanced Research. CD acknowledges support from TÜBİTAK (Project No:120Z594).

## AUTHORS’ CONTRIBUTIONS

C.D.T.C. contributed to Investigation, Formal analysis, Writing-Original draft preparation, Ç.D. contributed to Investigation, Conceptualization, Methodology, Supervision, Funding acquisition, Writing-Original draft preparation, Formal analysis, Writing™Reviewing and Editing

